# The appressorium of *Magnaporthe oryzae* remains mitotically active during post-penetration hyphal growth in rice cells

**DOI:** 10.1101/077669

**Authors:** Kiersun Jones, Cory B. Jenkinson, Jie Zhu, Sara Dorhmi, Chang Hyun Khang

## Abstract

To investigate the mitotic dynamics of an appressorium, we used time-lapse confocal imaging of a fluorescence-based mitotic reporter strain of *Magnaporthe oryzae*. We present evidence that: (i) appressoria remain viable and mitotically active after host penetration, (ii) appressorial mitosis, like invasive hyphal mitosis, is semi-closed, (iii) sister chromatids separate within the appressorium, (iv) a mitotic appressorial nucleus undergoes extreme constriction and elongation as it migrates through the penetration peg in a manner analogous to mitosis during cell-to-cell movement of invasive hyphae. These results provide new insight into the potential roles of the appressorium after host penetration and highlight the unique mitotic dynamics during rice blast infection.

## 1. Introduction

Many fungi form a specialized infection structure, called an appressorium, to directly penetrate into plant hosts (Howard and Valent, 1996; Ryder and Talbot, 2015). Recent studies have shown that the appressorium is considerably more complex than previously thought. It serves as the site of initial secretion of fungal effectors, and its development requires highly orchestrated events such as cytoskeleton reorganization and cell cycle regulation (Dagdas et al., 2012; Kleemann et al., 2012; Martin-Urdiroz et al., 2015; Ryder and Talbot, 2015; Saunders et al., 2010). There are questions yet to be answered, including how long the appressorium remains viable after host penetration, what mechanisms control division and migration of an appressorial nucleus, and whether the appressorium plays additional roles during post-penetration stages of infection. Here, we provide insights into some of these questions by studying the economically important rice blast disease caused by *Magnaporthe oryzae* (Khang and Valent, 2010). On the host surface, *M. oryzae* produces a single-celled appressorium, from which develops a penetration peg for breaching the plant surface. After penetration, the peg expands to form a filamentous invasive hypha (IH), which subsequently differentiates into bulbous IH. The appressorium provides a nucleus to the first IH cell, which continues to grow and divide for about 12 hours in the first-invaded cell, and then IH move into adjacent cells via IH pegs (Kankanala et al., 2007; Veneault-Fourrey et al., 2006). Recently, Jones et al (2016a) developed a fluorescence-based mitotic reporter strain of *M. oryzae* and provided evidence that IH undergo semi-closed mitosis and that a mitotic nucleus shows an extreme constriction and elongation when migrating through the IH peg to nucleate IH growing in an adjacent host cell. In this study, using the mitotic reporter strain coupled with high tempo-spatial resolution imaging, we show that the appressorium remains mitotically active during IH proliferation and that the appressorial nucleus undergoes extreme constriction and elongation through the penetration peg.

## 2. Results and discussion

### 2.1. The M. oryzae appressorium remains viable and mitotically active after host penetration

During live-cell imaging of rice cells infected with an *M. oryzae* strain expressing histone H1-mRFP and cytoplasmic EYFP, we unexpectedly observed that the appressorial cytoplasm and nucleus remained fluorescent even after IH had already spread into adjacent cells (Fig. 1A). Interestingly, we also observed two nuclei in an appressorium at the similar infection stage (Fig. 1B). These observations led us to hypothesize that the *M. oryzae* appressorium remains viable and even mitotically active while IH proliferate inside host cells. Upon further investigation, we found, consistent with this hypothesis, that 89.2% appressoria contained one or two fluorescent nuclei at 10 to 27-nuclear IH stage during first host cell colonization: of the 379 appressoria, 226 had one nucleus (59.6%), 112 had two nuclei (29.6%), and 41 had no nucleus (10.8%). Taken together, we propose that most appressoria retain viability and mitotic potential for many hours after host penetration.

**Figure 1.**
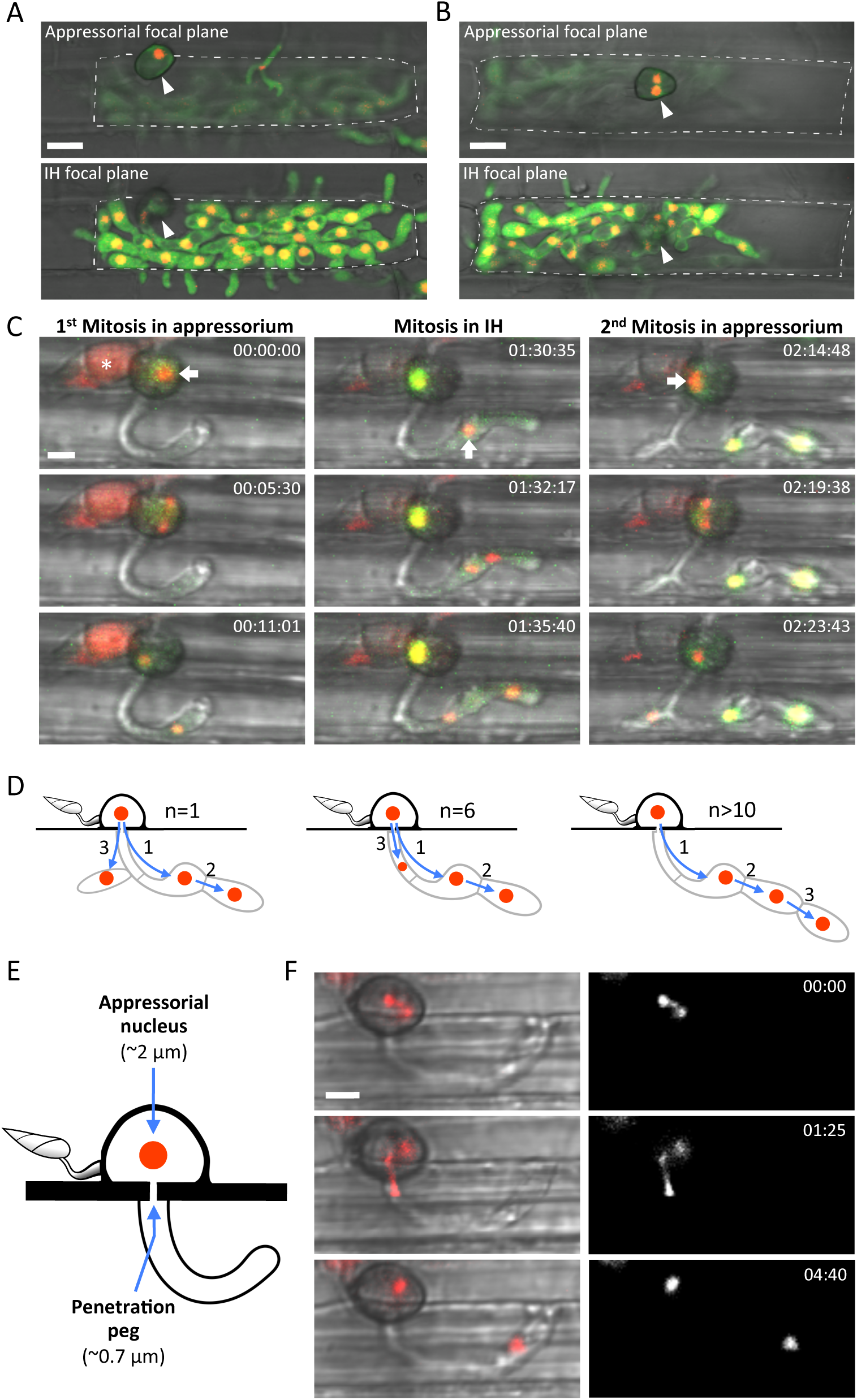
Appressorial mitotic dynamics of *M. oryzae* during invasion of rice cells. (A) Confocal images showing two focal planes of the same *M. oryzae* strain CKF110 infection at 36 hours post inoculation (hpi). This *M. oryzae* strain expresses histone H1-mRFP (red) and cytoplasmic EYFP (green). The IH focal plane (bottom) is 7 µm below the appressorial focal plane (top). A single nucleus in the appressorium (white arrowhead) is clearly visible in the appressorial focal plane. The IH focal plane shows highly branched IH that have spread into adjoining host rice cells. Bar = 10 µm. (B) Another infection at the same growth stage as (A). The IH focal plane (bottom) is 8 µm below the appressorial focal plane (top). Note that there are two nuclei in the appressorium (white arrowhead). Bar = 10 µm. (C) Time-lapse confocal images selected from Video 1, showing three rounds of mitosis during initial host colonization by *M. oryzae* CKF1962, starting at 25 hpi. This strain expresses histone H1-tdTomato (red; nuclear localization throughout the cell cycle) and GFP-NLS (green; nuclear localization during interphase but cytoplasmic localization during mitosis). The first mitosis is appressorial (left column), followed by mitosis in IH (middle column), and finally another appressorial mitosis (right column). Nuclei entering mitosis are denoted with white arrows. The conidium (white asterisk) is undergoing autophagic cell death. Times are shown in hours:minutes:seconds. Bar = 5 µm. (D) Schematic representations of the observed sequences of the first three rounds of mitosis after host penetration. Red circles indicate nuclei. Numbers associated with arrows indicate the order of nuclear divisions. Arrows indicate the direction of nuclear migration. The diagram on the left represents the mitotic sequence of the infection site in Fig. 1C and Video 1. The other sequences (middle and right) are of independently observed infection sites. (E) chematic diagram illustrating the question of how an appressorial nucleus (~2 µm in diameter) can migrate through the narrow penetration peg (~0.7 µm in diameter) to enter the filamentous IH. (F) Time-lapse single plane confocal images of *M. oryzae* CKF1962 invading a rice cell. Left: merged tdTomato and bright-field; right: tdTomato alone (shown in white). Sequence shows the first appressorial nuclear division and migration supplying a nucleus to the filamentous IH. The middle panel shows an extreme elongation and constriction of a mitotic nucleus as it migrates through the penetration pore. Times are shown in minutes:seconds. Bar = 5 µm.

### 2.2. Appressorial nuclear division and behavior

To investigate the dynamics of the appressorial nucleus, we used time-lapse confocal microscopy of the *M. oryzae* mitotic reporter strain CKF1962 infecting rice cells. This strain expresses histone H1-tdTomato, which remains associated with DNA throughout the cell cycle, together with GFP-NLS (nuclear localization signal), which localizes in the nucleus during interphase but disperses into the cytoplasm during semi-closed mitosis (Fig. 1C; Jones et al., 2016a). Dispersal of GFP-NLS from a nucleus occurs upon mitotic entry, thus allowing prediction of subsequent nuclear division and migration (Jones et al., 2016a; Shen et al., 2014). By taking advantage of this predictive capacity, we were able to capture nuclear division and migration in action (Video 1; Fig. 1C; Fig. 1F). We first identified an appressorium that had produced an anucleate filamentous IH and exhibited dispersed GFP-NLS with a single tdTomato-tagged nucleus, indicating the appressorium had just entered mitosis. We then began high temporal and spatial resolution confocal imaging (images taken every ~45 s during mitosis; 1 µm z-sections over 20 µm) (Video 1; Fig. 1C). The tdTomato-tagged sister chromatids separated within the appressorium, and then one mitotic nucleus moved into the developing first bulbous IH cell. The other nucleus remained in the appressorium (Fig. 1C left). Subsequently, we observed the growth and mitosis of the first bulbous IH cell (Fig. 1C middle). As we continued to follow the same infection site, we noticed an IH branching from the filamentous IH and then unexpectedly, another round of appressorial mitosis (Fig. 1C right). Subsequently, one nucleus migrated through the filamentous IH into the IH branch (Fig. 1C right; Fig. 1D left). This is the first demonstration of two rounds of appressorial mitosis supplying two nuclei to IH. We further found that the destination of the migrating nucleus can vary during 2^nd^ round appressorial mitosis. That is, the mitotic nucleus moved into the filamentous IH and stayed there if a new IH branch had not grown from it (n=6; Fig. 1D middle). In addition, appressorial mitosis did not always occur after the first round of IH mitosis. We often observed two sequential rounds of IH mitosis without the occurrence of a 2^nd^ appressorial mitosis (n>10; Fig. 1D right). Considered together, the *M. oryzae* appressorium can either undergo a second round of mitosis after the first round of IH mitosis or delay mitosis until a later IH growth stage.

### 2.3. Nuclear elongation and constriction through the penetration peg

The penetration peg is the narrow structure that pierces the host surface and conveys the appressorial contents, including the nucleus, into the growing IH (Howard and Valent, 1996). This raises an intriguing question of how the appressorial nucleus (~2 µm in diameter) moves through the constricted penetration peg (~0.7 µm in diameter; Howard and Valent, 1996) (Fig. 1E). Our high tempo-spatial resolution confocal imaging of the mitotic reporter strain revealed an extreme elongation (up to 13 µm) and constriction of the mitotic nucleus during migration through the penetration peg, followed by reformation into a typical spherical nucleus after entering the forming IH (Fig. 1F; n=8). The elongation and constriction of the appressorial nucleus was strikingly similar to previously reported nuclear dynamics during the spread of IH into adjacent host cells, in which the IH nucleus (~2 µm in diameter) constricted and elongated through the IH peg (~0.5 µm in diameter) at the host cell wall (Jones et al., 2016a). It is interesting to note that these nuclear dynamics occur during critical infection stages when the fungus gains the initial entry into the host cell using the appressorium-derived penetration peg and when it spreads into adjacent cells using the IH peg to continue intracellular colonization. The difference in size between both of the pegs and the nuclei likely requires specialized nuclear migration machinery to facilitate such extreme constriction and elongation of one mitotic nucleus while preventing the other nucleus from leaving the mother cell. The mechanisms controlling these unique mitotic dynamics remain to be determined.

In conclusion, we provide evidence that the appressorium remains viable and mitotically active well after host penetration. These observations expand our knowledge of the potential roles of the appressorium after nucleation of the first IH cell. The appressorium, in addition to being the critical penetration structure and the site of initial secretion of effectors, can also actively contribute additional nuclear material for IH growth beyond the first appressorial mitosis and likely continue to express effectors and other virulence factors. It remains to be determined whether persistence of the appressorium and retention of its mitotic potential significantly contribute to post-penetration proliferation. Our live-cell imaging has also revealed *M. oryzae*-specific cellular processes, i.e., nuclear constriction and elongation as well as semi-closed mitosis, which are very unlikely to occur in host plant cells. Understanding mechanisms underlying these pathogen-specific processes may provide new targets for disease control.

## 3. Methods

The mitotic reporter strain *M. oryzae* CKF1962 was previously described (Jones et al., 2016a). *M. oryzae* CKF110 was generated by transforming *M. oryzae* KV1 (Kankanala et al., 2007) with binary vector pBV217 using Agrobacterium-mediated transformation (Khang et al., 2005). pBV217 was produced by cloning of hH1 (histone H1 from *Neurospora crassa*) at the N terminus of mRFP under control of the constitutive promoter from the *M. oryzae* ribosomal protein 27 gene in binary vector pBHt2 (Mullins et al., 2001). Rice strain YT16 was grown and inoculated as previously described (Jones et al., 2016b).

Confocal microscopy was performed on a Zeiss Axio Imager M1 microscope equipped with a Zeiss LSM 510 META system using Plan-Apochromat 20X/0.8 NA and Plan-Neofluor 40x/1.3 NA (oil) objectives. Excitation/emission wavelengths were 488 nm/505 to 530 nm (EGFP and EYFP), and 543 nm/560 to 615 nm (tdTomato and mRFP). Images were acquired using LSM 510 software (Version 3.2). Video 1 was created with merged bright-field and fluorescence channels from informative focal planes selected from each of the 63 original confocal images (0.5-1.0 µm z-sections, covering 20-32 µm total) from the original time-lapse series. When documenting mitosis, we acquired z-stack images once every ~45 s. Upon completion of GFP reimport into a nucleus we reduced the image acquisition rate to once every few minutes (up to 23 minutes) in order to minimize potential phototoxicity while continuing to document IH growth. Images were analyzed and processed using a combination of the Zen software, Adobe Photoshop, and ImageJ (http://imagej.nih.gov/ij/).

## Acknowledgements

We thank all members of the Khang Lab (http://www.khanglab.org/) for their help and discussion. We acknowledge the assistance of the Biomedical Microscopy Core at the University of Georgia with imaging using a Zeiss LSM 510 confocal microscope. This work was supported by the Agriculture and Food Research Initiative competitive grants program, Award number 2014-67013-21717 from the USDA National Institute of Food and Agriculture.

---

**Video 1.**

Time-lapse series showing the first three rounds of nuclear division in *M. oryzae* CKF1962 after host penetration (beginning at 25 hpi). The first mitosis occurs in the appressorium, followed by IH growth and mitosis in IH. Then, an IH branches from the filamentous IH and becomes nucleated upon a second round of appressorial mitosis. Nuclear positions are revealed by histone H1-tdTomato (red) throughout the cell cycle, while GFP-NLS (green) associates with nuclei only during interphase and disperses into the cytoplasm during mitosis. Times are shown in (hr:min:sec). Bar = 10 µm.

